# Epigenetic Regulation During Flatfish Metamorphosis: Integrative Omics Analysis of DNA Methylation and Gene Expression in Turbot Brain

**DOI:** 10.1101/2025.02.20.639312

**Authors:** Laura Guerrero-Peña, Paula Suarez-Bregua, Núria Sánchez-Baizán, Francesc Piferrer, Juan J. Tena, Josep Rotllant

## Abstract

The regulation of gene expression plays a pivotal role in the complex metamorphic process of flatfishes, which oversees the dynamic alterations that occur during their transition from a pelagic to a benthic way of life. Flatfish metamorphosis is characterized by substantial modifications in both form and function, which are initiated by thyroid hormones. Epigenetic mechanisms play a vital role in the regulation of gene expression during this transformative process. This study examines the molecular mechanisms underlying flatfish metamorphosis by integrating multi-omics data that provide information on chromatin status, DNA methylation profile, and gene expression at three key stages (pre-metamorphosis, climax and post-metamorphosis) in the turbot (*Scophthalmus maximus*) brain, a complex organ that plays a critical role in the regulation of this intricate process. The analysis of DNA methylation patterns revealed major epigenetic dynamic changes during the different metamorphosis stages. Specifically, the methylation levels exhibited a typical bimodal distribution in the pre-metamorphic stage, transitioned to intermediate levels of methylation (around 50%) during the climax stage, and reverted to a bimodal distribution in the post-metamorphic stage. Notably, DMRs were identified in regions of open chromatin that colocalized with CpG islands. Moreover, our results indicate an inverse relationship between DNA methylation and transcriptional activity in regions near the transcription start sites (TSSs), where high levels of methylation correspond to low expression and low levels of methylation corresponds to high gene transcription. This study thereby elucidates the regulatory impact of methylation on gene expression during the metamorphosis process in the flatfish brain.

## Introduction

Flatfish (Pleuronectiforms) exhibit the capacity to adapt to the benthic environment despite being born with a morphologically symmetrical anatomy that is adapted to the pelagic environment. In the early stages of their development, flatfish undergo extensisve phenotypic changes that result in a completely asymmetrical cranial structure (***Okada et al., 2001***; ***Brewster, 1987***; ***Schreiber, 2006***), with both eyes located on one side of the body (***Sæle et al., 2006***; ***Sæle***, ***Øystein; Smáradóttir, Heiodís; Pittman, 2006***), and behaviour and mobility adapted to the seafloor (***Gibson et al., 2015***). These changes are coordinated and promoted by thyroid hormones (***de Jesus et al., 1993***; ***Gomes et al., 2015***; ***Deal and Volkoff, 2020***), which trigger metamorphosis in flatfish. Despite the acknowledged importance of thyroid hormones in metamorphosis, the coordinated regulation of the molecular processes that ensure the metamorphic remodelling remains unknown.

Among these multiple molecular processes, gene regulation plays a fundamental role, orchestrating the dynamic transformations that occur during the transition from one life stage to another (***Raj et al., 2023***; ***Alves et al., 2016***). During the metamorphosis, flatfish undergo profound morphological and physiological changes. The intricate interplay of epigenetic mechanisms, including chromatin modification such as DNA methylation (***Wu et al., 2018***; ***Covelo-Soto et al., 2015***) and histone modification (***Bannister and Kouzarides, 2011***), and chromatin remodelling (***Wolffe and Hayes, 1999***; ***Chen and Dent, 2014***) contribute to the tight regulation of gene expression patterns. This precise molecular orchestration ensures the successful progression of developmental programs (***Bogdanović et al., 2011, 2017***), ultimately leading to the determination of the juvenile lifestyle and body plan.

Open chromatin, which is indicative of accessible genomic regions, facilitates the interaction of transcription factors and regulatory machinery with DNA (***Klemm et al., 2019***; ***Chai et al., 2013***). This allows the activation or repression of gene expression (***Wu, 1997***; ***Meier and Brehm, 2014***). Chromatin dynamics and DNA methylation are closely related (***Huck-Hui and Bird, 1999***). Consequently, when DNA is in a methylated state, nearby histones become deacetylated, resulting in compact and temporarily silent chromatin. DNA cytosine methylation represents a pivotal epigenetic mechanism with evident implications for gene silencing, particularly within a CpG context in vertebrates (***Lyko, 2017***). Vertebrate genomes exhibit high levels of CpG methylation, with a fraction of unmethylated regions coinciding with CpG-rich regions, designated as CpG islands (CGI). These regions are predominantly localized on gene promoters. Prior research has identified substantial alterations in DNA methylation at pivotal stages of metamorphosis in *Xenopus*, underscoring the critical role of methylation in the gene regulatory network during this process (***Kyono et al., 2020***). Nevertheless, despite the extensive research on metamorphosis in diverse flatfish species, the epigenetic regulation of flatfish metamorphosis remains relatively understudied, particularly with regard to the interplay between epigenetics and transcriptional activity.

It is therefore imperative to gain information of the landscape of chromatin methylation and structure in order to understand its role in the regulation of gene expression. In this study, we have investigated the molecular processes underlaying the metamorphosis. Using multi-omics data, we integrated information on chromatin status, DNA methylation profile, and gene expression at three key stages (pre-metamorphosis, metamorphic climax and post-metamorphosis) of metamorphic development of the turbot (*Scophthalmus maximus*) brain, a complex organ that plays a pivotal role in regulating this process. We identified differentially methylated regions (DMRs) in the accessible chromatin at the climax of metamorphosis, with a significant proportion of these regions colocalizing with CpG islands. Furthermore, our data also reveal unique regions with an intermediate methylation levels (%mCG/CG around 50) at the climax of metamorphosis. Finally, this study confirms the inverse correlation between DNA methylation and gene expression in regions close to the TSSs in turbot brain.

## Results

### Disparities in DNA methylation during the metamorphic developmental stages of flatfish

In order to gain insight into the regulatory mechanisms underlying metamorphosis in flatfish, genomewide methylation profiles were compared at three key stages of turbot development: pre-metamorphic, metamorphic climax and post-metamorphic stages. The mean number single-end reads per brain sample was 24 million, with an average alignment rate of 91.87% to the most recent turbot genome. Principal component analysis demonstrated that the pre-metamorphic and climax stages exhibited reduced stability and consistency among samples in comparison to the post-metamorphic stage ***Figure 1a***. However, no significant differences in overall methylation levels were observed between stages ***Figure 1b***. Approximately 50% of CpGs in all our samples exhibited high levels of methylation, while only about 30% of the CpGs in the genome were unmethylated.

**Figure 1.**
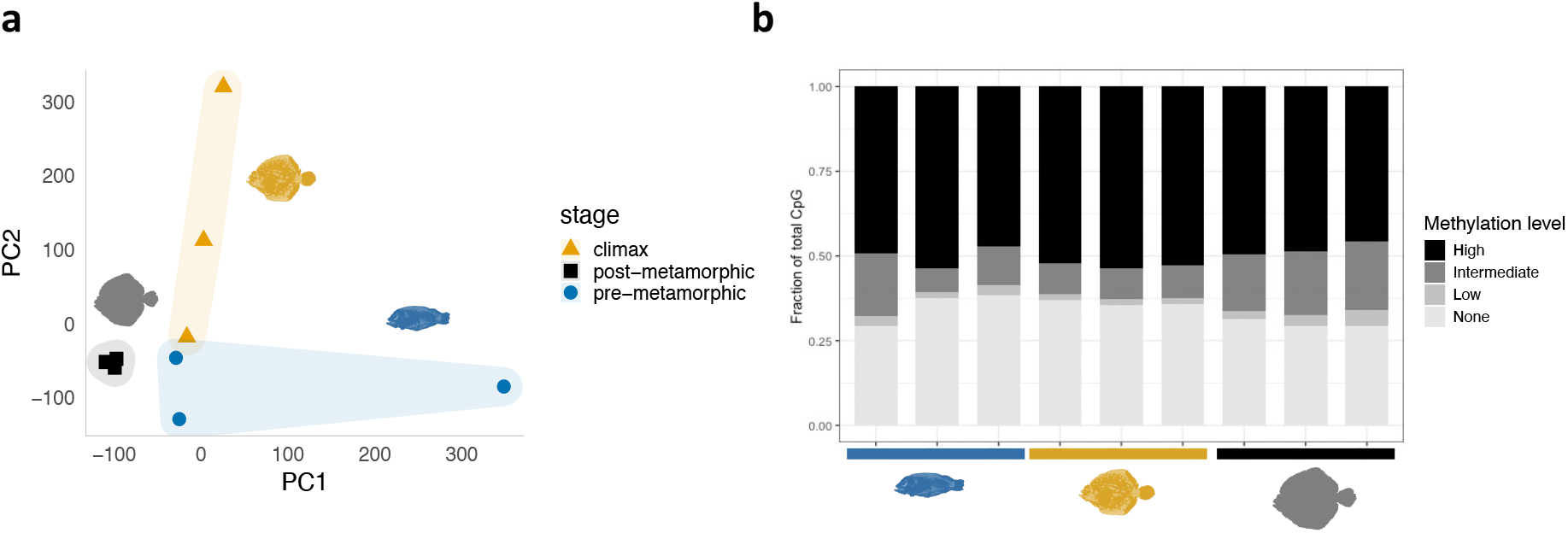
Figure 1. Methylome data from turbot brain at three different metamorphic stages. (a) Principal component analysis (PCA) plot illustrating the clustering pattern of samples based on DNA methylation data using around 250.000 CpGs. PCA1: 15.8% variance; PC2: 15.3% variance. Each point represents an individual sample. (b) Global DNA methylation levels per metamorphic stage and across all samples, classified according to the following categories: High methylation 0.8-1; Intermediate methylation > 0.2-0.8; Low methylation > 0-0.2; and No methylation = 0. The silhouettes of fish represent the following stages: pre-metamorphic in blue, climax of metamorphosis in yellow and post-metamorphic in black.

A pairwise comparison analysis was conducted to identify differentially methylated regions (DMRs) between developmental stages: 856 DMRs between the pre-metamorphic and climax stages, 10626 DMRs between climax and post-metamorphic stages, and 1816 DMRs between the pre-metamorphic and postmetamorphic stages***Figure 2a***, with a mean DMR length of 1.8 kb. It is noteworthy that an examination of the mCG/CG of the DMRs revealed that the pre-metamorphic and post-metamorphic stages exhibited a bimodal distribution, with regions having either high or low methylation levels. In contrast, at the climax of metamorphosis exhibited a unimodal distribution, with the majority of the regions displaying intermediate mCG/CG (around 50%) ***Figure 2b***. With respect to the methylation gain and loss across the developmental stages, the data indicate that methylation gain at the climax stage was observed to be 96.24% and 45.13% compared to pre-metamorphic and post-metamorphic stages, respectively. Although there is only a 4.86% gain in methylation between the pre-metamorphic stage and climax stage, the differences in mCG/CG did not exceed 21%. Similarly with the post-metamorphic stage, differences of up to 26% were observed. DMRs were primarily enriched in promoter and exon regions, while introns and intergenic regions exhibited a lower representation. This difference was especially evident in the contrast of pre-metamorphosis and metamorphosis climax ***Figure 2c***. The GO enrichment analysis revealed that the development and differentiation process were the most prominent pathways associated with DMRs when comparing the stages ***Figure 2d***.

**Figure 2.**
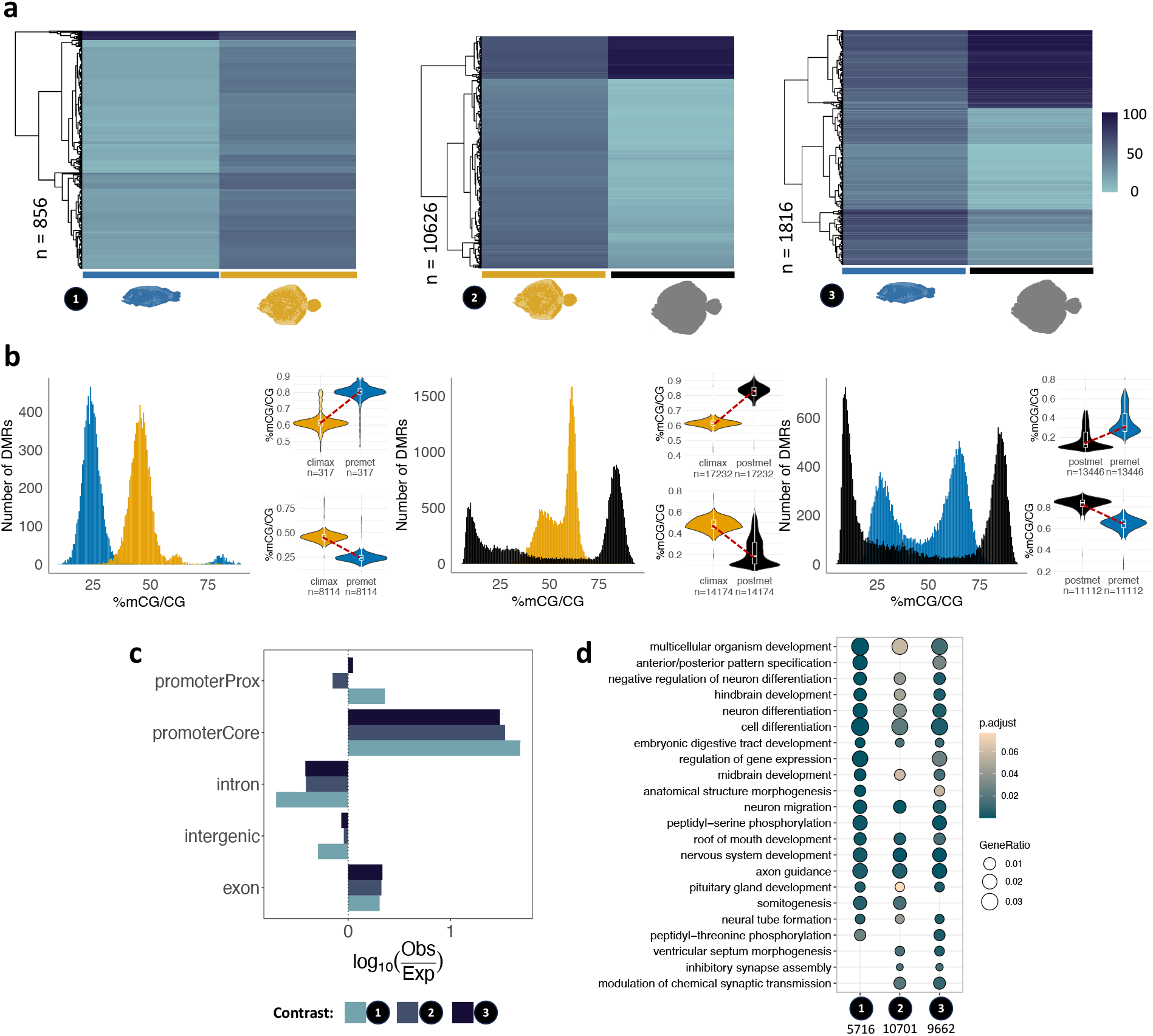
Figure 2. DNA methylation profile throughout turbot metamorphosis performing pairwise comparisons among stages. (a) Heatmaps illustrating the percentage of mCG/CG in DMRs identified across metamorphic stages. The colour intensity represents the average percentage of mCG/CG within each DMR. The number of DMRs identified for each comparison is indicated on the left-hand side of each heatmap. (b) A bar plot illustrating the distribution of mCG/CG among the identified DMRs for each contrast. The x-axis represents the percentage of mCG/CG, while the y-axis denotes the number of DMRs. Violin plots of the gain and loss of methylation with the number of DMRs indicated below. The percentage of mCG/CG in DMRs is represented for each metamorphic stage: blue, pre-metamorphic; yellow, climax of metamorphosis; and black, post-metamorphosis stage. (c) The distribution of DMRs across genomic features in comparison to the expected distribution. The sizes of the PromoterProx and PromoterCore regions are located 2000 and 100 base pairs upstream from the transcription start site (TSS), respectively. (d) Gene ontology (GO) enrichment analysis of genes associated with DMRs. The number of genes that are enriched in each comparison is provided below the plot. The colour of the dots indicates the p.adjusted value, while their size represents the gene ratio. The silhouettes of fish are represented by the following colours: blue for the pre-metamorphic stage, yellow for the climax of metamorphosis and black for the post-metamorphic stage. The numbers in black indicate the results of the pairwise comparisons: 1, pre-metamorphic versus climax stage; 2, climax versus post-metamorphic stage; and 3, pre-metamorphic versus post-metamorphic stage.

### Chromatin status of differentially methylated regions

Methylation can be observed in both accessible chromatin regions and in compacted and inactive regions. Each context has different implications for the gene regulatory network. The initial analysis was conducted using an outside-in approach, focusing on stage-specific DMRs, including also DMRs with a methylation difference of less than 25% between stages. A single dataset was created to encompass all identified DMRs, which revealed three distinct clusters ***Figure 3a***. A total of 42169 DMRs were identified across the stages. Unexpectedly, over 80% of DMRs (26160 DMRs) were hypermethylated exclusively at the climax stage of metamorphosis. A limited number of DMRs exhibit exclusive hypermethylation at the pre-metamorphic (1729 DMRs) and post-metamorphic (2499 DMRs) stages. Furthermore, we observed a substantial number of DMRs exhibiting subtle methylation levels between the pre-metamorphic and climax stages. Focusing exclusively on the hypermethylated DMRs at the climax of metamorphosis, we integrated DNA methylation data of specific regions with chromatin accessibility data, using ATAC-seq data (GSE215396) (***Guerrero-Peña et al., 2023b***) from whole fish at four different stages of metamorphosis (phylotypic, pre-metamorphic, climax and post-metamorphic stages). The results revealed that of the exclusively hypermethylated DNA methylation regions (DMRs) at the climax stage ***Figure 3a***, 84% were situated in a closed or non-accessible chromatin region ***Figure 3b***, while the remaining 16% were located in an open chromatin region (OCR) ***Figure 3b***. It is noteworthy that the aforementioned correlation between methylation ***Figure 3a*** and chromatin status ***Figure 3b*** observed in the climax stage is also evident in the premetamorphic stage, revealing a developmental conserved pattern.

**Figure 3.**
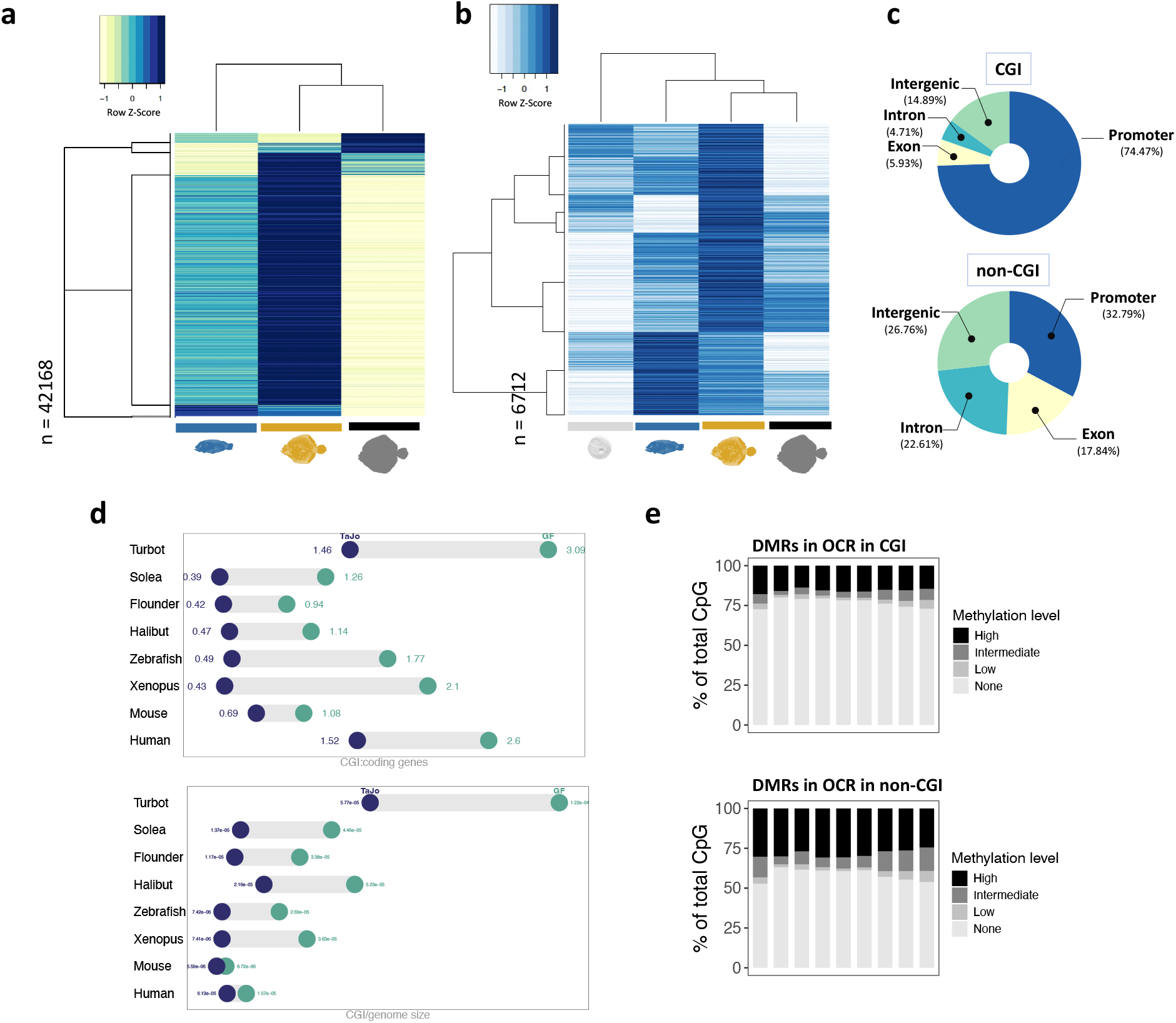
Figure 3. Analysis of methylated and open chromatin regions at metamorphosis stages of turbot. (a) Hierarchical clustering of the differentially methylated regions (DMRs) identified among metamorphosis stages displaying an open chromatin region in at least one of the three key stages. (b) A hierarchical clustering analysis was conducted using open peak regions that overlap with DMRs that were uniquely hypermethylated during the climax of metamorphosis. (c) Genomic features overlapping with the dataset of DMRs and in OCRs at the climax of metamorphosis categorized into distinct genomic regions. (d) Ratio of CGI/coding genes and CGI/genome size in four species of flatfish and other model organisms. The identification of CGIs was performed using two distinct algorithms: the Takai and Jones algorithm is represented in blue, while the Garden-Gardiner and Frommer algorithm is represented in green. In (a) and (b) the silhouettes of fish are represented in white, blue, yellow and black to indicate the phylotypic, pre-metamorphic, climax of metamorphosis and post-metamorphic stages, respectively. (e) The ratio of methylcytosine to cytosine in differentially (mCG/CG) in DMR in OCR in CGI and in DMRs in OCR in non-CGI at the climax of metamorphosis. The categories are defined as follows: High methylation 0.8-1; Intermediate methylation > 0.2-0.8; Low methylation >0-0.2; and No methylation = 0.

### CpG islands annotation and overlap detection with DMRs in metamorphic stage

Subsequent to the outside-in analysis, we examined hypermethylated DMRs in climax and in an OCR at the same stage. As anticipated, the majority of DMRs in OCR in climax were situated in a CpG-rich regions (71.83%), also designated as a CpG islands (CGI). Furthermore, 78.01% of DMRs located in both OCR and in CGI at climax were identified within promoter regions (upstream = 5000; downstream = 100 of TSS) ***Figure 3c***.

CpG islands (CGIs) possess distinctive characteristics, as they represent the unmethylated portion of the genome and are typically situated within an open chromatin region (OCR). The number of CGIs in vertebrates exhibits considerable variability (***Bird, 1980***). In order to gain a broader understanding of the subject matter, we identified and compared CGIs in other organisms, including humans, mice, zebrafish, and other flatfish species. To ensure accuracy, two different CGI identification algorithms were used, including the methods of ***Takai and Jones (2002***) and the algorithms of ***Gardiner-Garden and Frommer (1987***). Unexpectedly, the number of CGIs in turbot was considerably higher than in other model species. Moreover, these differences were maintained when compared to other flatfish species, exhibiting a distinctive proportion of CGIs in the genome ***Figure 3d***. We compared the overall methylation of DMRs in regions that colocalized with CGI to those that did not. As anticipated, DMRs that colocalized with CGI demonstrated significantly lower methylation levels ***Figure 3e***. This validates the reliability of our data and the algorithms utilized for CGI identification.

### The correlation between methylation in accessible chromatin regions and gene expression

Despite the association between DNA methylation and gene silencing, the role in facilitating transcriptional changes during metamorphic processes remains unclear. The objective of this study was to investigate whether there is a correlation between transcriptional activity and DNA methylation marks in across the turbot genome. RNA sequences from three developmental stages were aligned to the reference genome, reaching and average alignment of 96%. Analysis of gene expression profiles throughout metamorphosis showed stage-specific expression patterns ***Figure 4a***, revealing that, in contrast to the methylation profile, the lowest number of exclusively overexpressed genes were in the metamorphic climax. This divergent pattern between methylome and transcriptome suggests a link between DNA methylation and gene silencing in turbot. We identified 284, 39 and 1165 unique overexpressed genes in the pre-metamorphic, climax of metamorphosis and post-metamorphic stages, respectively. The expression levels were classified into 10 expression deciles, with decile 1 representing the lowest and 10 the highest.

**Figure 4.**
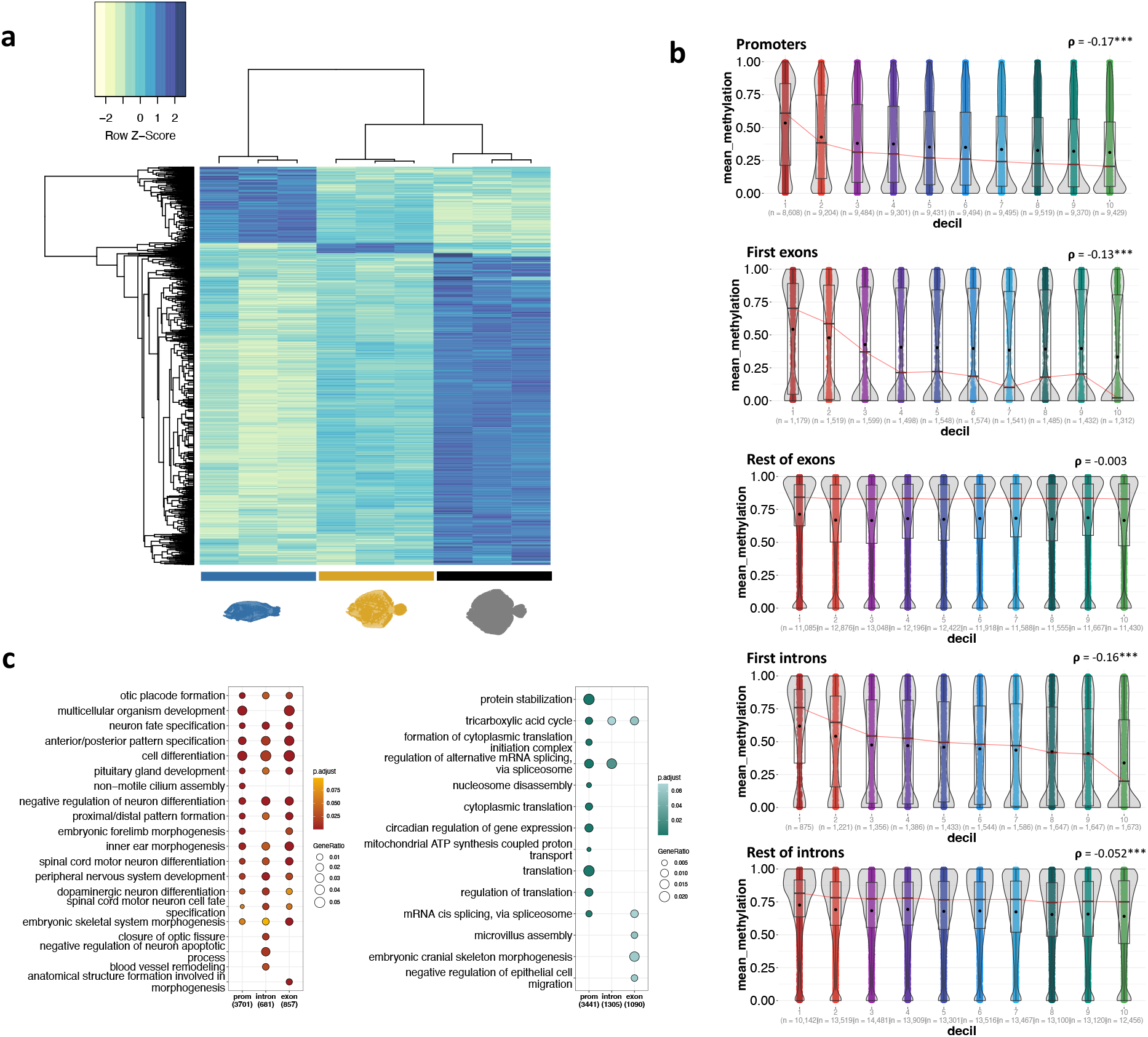
Figure 4. Correlation between DNA methylation and gene expression in the turbot brain during metamorphosis. (a) Hierarchical clustering of the gene expression matrix demonstrates a set of genes that are uniquely overexpressed at each stage of metamorphosis. (b) The Spearman correlation between the percentage of mCG/CG and the normalized gene expression levels, which have been categorised into deciles, was evaluated in the following regions of the turbot genome: promoter, first exon, the rest of exons, first intron and the rest of introns. The distribution of gene expression for each decile is illustrated by violin plots. The boxplot shows the interquartile range (Q1-Q3), with the median represented by a line and mean by a dot. The results are reported with Spearman’s rank correlation coefficient (ρ) and the associated significance levels (*** = p<0.001; * = p<0.05). (c) Gene ontology (GO) enrichment of the three highest and three lowest gene expression deciles for the promoter, first exons and first introns. The silhouettes of fish in blue, yellow and black represent the pre-metamorphic, climax of metamorphosis and post-metamorphic stages, respectively.

This study also explores the methylation levels associated with different gene regions. A negative correlation was observed between DNA methylation and gene expression in the promoter (Spearman’s rank correlation ρ = -0.17, P<2.2e-16), first exon (Spearman’s rank correlation, ρ = -0.13, P<2.2e-16) and first intron (ρ = -0.16, P<2.2e-16). Nevertheless, no correlation was discernible in the remaining exons and introns ***Figure 4b***. The first three deciles, indicative of low expression, correlated with high methylation levels, while the final three deciles, representative of high expression, correlated with very low methylation levels across all samples. We found 8608 genes in three first deciles while 10880 genes were identified in the last three deciles. Of these, 79.44% and 76.66%, respectively, were found to be located in promoters, supporting the idea that the regulation of gene expression is predominantly concentrated in promoters, particularly in the context of repression. To elucidate the functions regulated by methylation of each region (promoter, first exon and first intron), we conducted a Gene Ontology (GOs) analysis on genes exhibiting highly or low expression levels that correlated with DNA methylation. Methylation-repressed genes were enriched for molecular functions related to morphogenesis. In contrast, highly expressed methylation-associated genes were enriched for processes related to chromatin organization, translation and RNA splicing. Furthermore, we identified functions that are specifically regulated by methylation in the first exons, including microvillus assembly, embryonic cranial skeleton morphogenesis and negative regulation of epithelial cell migration ***Figure 4c***. These findings suggest that during metamorphosis in flatfish chromatin is highly dynamic, and that post-transcriptional mRNA and translational processes are highly active. This indicate that this is a perfectly synchronized process regulated by different mechanisms.

## Discussion

Metamorphosis is a biological process by which an animal undergoes physical and physiological development. This process is characterized by a remarkable and relatively rapid change in the animal’s body structure. The hypothalamus-pituitary-thyroid (HPT) axis regulates the production and release of thyroid hormones (TH) (***Deal and Volkoff, 2020***; ***Power et al., 2008***), essential for initiating and controlling the dramatic physical changes that occur during metamorphosis (***de Jesus et al., 1993***; ***Miwa and Inui, 1987***; ***Campinho et al., 2018***). Through this hormonal regulation, the brain influences cell growth and differentiation, leading to the transformation of the animal’s body structure.

Although it is well stablished that TH functions as an epigenetic switch, modifying chromatin structure and facilitating transcriptional activation (***Shi et al., 2012***; ***Cheng et al., 2010***), the alterations in DNA methylation that occur during the post-embryonic development of flatfish in the brain remains poorly understood. The objective was to examine the impact of epigenetic regulation on gene expression changes during the metamorphosis process of the flatfish brain. To this end, global brain methylation was assessed in turbot at three key developmental stages: pre-metamorphic, metamorphic climax, and post-metamorphic. Despite the expectation of higher methylation levels in later stages, which would be in contrast with the significantly lower levels observed in the early stages (***Huidobro et al., 2013***), our study did not find such differences during turbot development. These findings also diverge from those of other studies, which have identified discrepancies in global methylation ratios during metamorphosis in turbot (***Suarez-Bregua et al., 2020***) and other organisms (***Covelo-Soto et al., 2015***; ***Trautner et al., 2017***; ***Suarez-Bregua et al., 2021***) using methylation-sensitive amplified polymorphism (MSAP) techniques. However, the observed trend of high CpG methylation levels (> 50%) across all turbot developmental stages aligns with the descriptions provided for the brain in other vertebrates (***de Mendoza et al., 2021***) and, more specifically, in actinopterygians (***Klughammer et al., 2023***). The conserved methylation levels observed across the developmental stages appears to be due to a balance of DNA methylation and demethylation processes. This is evidenced by the fact that the results of comparisons between stages do demonstrate significant differences in methylation in specific regions. It was surprising to find that all differentially methylated region (DMRs) identified in pairwise comparisons between stages showed an intermediate methylation pattern (mCpG/CpG \approx 50%) at the climax stage. However, a clear bimodal distribution was observed in these DMRs, exhibiting either full methylation or complete unmethylation at both the pre-metamorphic and post-metamorphic stages. During metamorphosis, there is a drastic change in environmental conditions. The intermediate methylation shown by DMRs in the climax stage may be attributed to the presence of epigenetically bistable regions, suggesting that only a subset of cells undergoes methylation changes in response to the demands to transition from larval to adult stage (***Brindley et al., 2022***). This results in the establishment of transcriptional “on” and “off” states that facilitate behavioural and neuronal adaptation, enabling the activation or repression of genes as required throughout the process. Although these intermediate methylated regions are often associated with loss of gene expression, the functional interpretation is unreliable (***Hay et al., 2023***). However, they may be involved in fine-tuning gene expression rather than serving as on/off switches. This fine-tunned regulation could be essential for the coordinated expression of genes involved in development, tissue differentiation, and morphological changes that occur during metamorphosis.

Moreover, our data showed a higher degree of methylation gain for both contrasts, namely, climax versus pre-metamorphic stages and climax versus post-metamorphic stages. It is expected that at later development stages, the genome will exhibit a greater number of hypermethylated DMRs. However, our data indicated that there were more demethylation processes from an intermediate methylation state when comparing the climax stage with the post-metamorphic stage. It appears that there is a continued need for regions that are capable of being malleable in response to external factors. Furthermore, DMRs were predominantly located in promoters and exons. The pre-metamorphic/climax contrast exhibited a notable low number of DMRs in intronic and intergenic regions than anticipated, aligning with DMRs of intermediate methylation gain for the climax. Exon methylation is linked to the inclusion of exons (***Shayevitch et al., 2018***; ***Elliott et al., 2015***), which regulate alternative splicing processes, potentially leading to enhanced protein diversity during metamorphosis and facilitating phenotypic plasticity in the organism.

A well-established correlation exists between DNA methylation and chromatin structure (***Huck-Hui and Bird, 1999***; ***Razin, 1998***). Our data demonstrate that among the DMRs exhibiting exclusive hypermethylation during the climax stage of metamorphosis, only 25% were identified in an accessible chromatin region within the same stage. This corroborates de notion that methylation not only regulates gene expression but also plays a role in maintaining chromatin in a compact and inaccessible conformation (***Cedar and Bergman, 2009***). Furthermore, DMRs in OCR at the climax colocalized with CGI in 70% of observations. CGIs represent the unmethylated portion of the genomeand are typically localized around promoters (***Larsen et al., 1992***). The high incidence of DMRs in OCR colocalising with CGI is to be expected, given that CGIs are often found in highly accessible regions of the genome. Indeed, it has been observed that nucleosomes assemble in an unstable manner in CGI regions (***Ramirez-Carrozzi et al., 2009***), with low levels of H1 and high acetylation of histones H3 and H4 (***Tazi and Bird, 1990***).

Despite extensive research on the CpG-rich regions of the genome and their influence on chromatin structure and transcriptional regulation, the definition of a CGI remains somewhat arbitrary. Thus, in this study, CGIs in the turbot genome were identified using two proposed algorithms: ***Gardiner-Garden and Frommer (1987***) (GGF) and ***Takai and Jones (2002***) (TJ). Although the GGF algorithm identified a significantly greater number of CGIs, this is likely due to the inclusion of repetitive regions (***Han and Zhao, 2008***). Nevertheless, the proportion of CGIs in comparation to other organisms remains consistent. Accordingly, the application of both the GGF and TJ algorithms revealed that the number of CGIs identified in turbot is markedly higher than in other organisms, and notably, also higher than in other flatfishes. As CGIs represents the unmethylated portion of the genome, they offer protection against deamination, the mutation of methylated CpG to TpG (***Bird, 1980***; ***Coulondre et al., 1978***). Nevertheless, the hypothesis that the loss of CGIs may be attributed to the novo methylation in CGIs has been proposed (***Antequera and Bird, 1993***; ***Jiang et al., 2007***). This hypothesis is supported by the observation that a greater number of CGIs have been identified in lamprey, which is a more ancestral organism than the others. However, this hypothesis does not appear to be valid for turbot, as the number of CGIs identified differs significantly despite the evolutionarily proximity of turbot to zebrafish and other flatfishes.

Our data indicate that a considerable number of DMRs were identified in OCR during turbot metamorphosis. A direct correlation between DNA methylation and gene silencing has been described in vertebrates, although not all regions of the genome respond equally to methylation (***Anastasiadi et al., 2018***). Our findings corroborate the notion of an inverse correlation between DNA methylation in promoters, first introns and first exons and gene expression. This supports the proposition that first introns possess distinctive regulatory properties with respect to the rest of the genome (***Li et al., 2012***; ***Jo et al., 2019***; ***Majewski and Ott, 2002***). However, our results did not yield evidence for the regulation of biological processes associated with specific genomic regions that respond to high levels of methylation.

To conclude, in this study we examined the regulatory processes of gene expression, specifically focusing on the roles of DNA methylation and chromatin structure during the metamorphosis of flatfish. The data demonstrate a consistent level of global methylation across all stages. However, an intermediate methylation profile was identified in regions with differential methylation during the climax of metamorphosis, which appeared to be a stage of adaptation to new conditions. Moreover, a considerable proportion of the differentially methylated regions exhibited hypermethylation at the climax stage, in comparison to the pre- and post-metamorphosis stages. Conversely, a notably high number of CpG-rich regions were identified in turbot in comparison to other species. Furthermore, our data supported the correlation between methylation in regions adjacent to the TSS and gene expression in turbot. Collectively, our dataset illustrates and corroborates the mechanisms that facilitate higher neuronal plasticity during metamorphic remodelling in flatfish.

## Methods and Materials

### Fish and sampling

Newborn turbot (*Scophthalmus maximus*) used in this study were raised under a typical commercial production system and provided by the company Insuiña S.L, Grupo Nueva Pescanova (Pontevedra, Spain). Fish from a single mating pair were collected at various stages by skilled company personnel. The number of fish used for all experimental procedures was determined based on the minimum number needed to achieve statistically reliable and robust results. The metamorphic stages were defined according to the morphological criteria described by Al-Maghazachi and Gibson (***Al-Maghazachi and Gibson, 1984***) and Suarez-Bregua (***Suarez-Bregua et al., 2020***), focusing on eye migration, overall body symmetry, and rearing temperature (21°C): pre-metamorphic stage (13 days post-fertilization, dpf]), characterized by symmetrical larvae before eye migration. Metamorphic climax (21 dpf), characterized by larvae showing asymmetrical features with the upper edge of the right eye visible from the left side. Post-metamorphic stage (44 dpf), characterized by asymmetrical juveniles with fully migrated eyes. Individual fish at each metamorphic stage (n = 3 independent biological replicates per stage) were euthanized using a lethal dose of MS-222 (Sigma-Aldrich, USA) (500mg/L for 30-40 minutes) and brains dissected with the aid of a Leica M165FC stereomicroscope equipped with a Leica DFC310FX camera (Leica, Germany). Ethical approval for the conduct of all studies (ES360570202002/17/FUN.1/BIOLAN.08/JRM) was obtained from the Institutional Animal Care and Use Committee of the IIM-CSIC Institute. This aligns with the guidelines set forth by the National Advisory Committee for Laboratory Animal Research, which was authorized by the Spanish Authority (RD53/2013). This research was conducted in accordance with the European animal directive (2010/63/UE), which is designed to protect experimental animals.

### DNA isolation and library preparation

Entire brains from turbot were carefully dissected, by first cutting off the head and then extracting the brain by opening the skull. All the samples were treated with DNAse-free RNAse (final concentration 2 µg/µl) and protease, to prevent interference from RNA and binding proteins, and were then subjected to mechanical homogenization. DNA extraction and purification were performed using the Blood & Cell Culture DNA kit (Qiagen) according to the manufacturer’s instructions. Extracted DNA was quantified using Qubit fluorometric quantitation (ThermoFisher Scientific) before being utilized for library construction.

Reduced representation bisulfite sequencing (RRBS) libraries were prepared from DNA extracted from 9 individual brain samples using Premium RRBS kit from Diagenode^®^, in strict accordance with the manufacturer’s guidelines. For each sample, a total of 100 ng of DNA was used for MspI digestion, which specifically targets CCGG sites. After digestion, 5’ CG overhangs were repaired, and A-tails were added. End repair was followed by adaptor ligation and quantification by qPCR. Sample pooling was performed based on the individual sample amounts determined by qPCR. The resulting DNA fragments were bisulfite converted, and the optimal number of enrichment amplification cycles was determined by qPCR. After clean-up step using AMPure XP Beads, concentrations were measured using the Qubit High Sensitivity Assay (ThermoFisher Scientific). The final RRBS libraries were sequenced on an Illumina HiSeq2000 platform, using 50 bp single end reads.

### DNA Methylation Analysis and Differential Methylated Regions Prediction

The quality of sequenced data was evaluated using FastQC v0.12.0 tool, and adapters were removed using the TrimGalore v06.10-0 (***Krueger et al., 2023***) software. The efficiency CpG bisulphite conversion was verified through the analysis of a spike-in control. Subsequently, the RRBS sequenced reads were aligned to the turbot reference genome (ASM1334776v1) with BSMAP v2.90 (***Xi and Li, 2009***), and methylation was performed using the *methratio*.*py* Python script, included in the BSMAP software. A differential methylation analysis was conducted among the preselected stages using the DSS v2.50.1 (***Wu et al., 2015***) R package. Pairwise comparisons between developmental stages (pre-metamorphic, climax and post-metamorphic stages) were conducted. To detect differential methylation, statistical tests were performed at each CpG site and based on a rigorous Wald test for beta-binomial distributions using the DMLtest function (smoothing = TRUE). The statistical test considers biological variations between replicates and the depth of sequencing. Differentially methylated regions (DMRs) were defined as regions that met the following three criteria: (1) p.threshold \leq 0.05; (2) minCG = 10; (3) mean methylation differences \geq 25%.

### Total RNA Isolation and Sequencing

RNA was extracted from each brain, with three independent replicates per stage. The RNA was subsequently purified using the RNeasy Mini Kit (Qiagen) in accordance with the manufacturer’s protocol. RNA integrity was evaluated using an Agilent 2100 bioanalyzer (Agilent Technologies, USA), and only samples with an RNA integrity number (RIN) equal to or exceeding 8 were included in subsequent analyses. Strand-specific mRNA libraries were sequenced on a DNBseqTM platform, generating paired-end reads of 100 base pairs (bp) length for each sample.

### RNA sequence alignment and differential expression analysis

High-quality RNA-seq raw reads were aligned to the turbot genome assembly (ASM318616v1) using the STAR v2.7.0e (***Dobin et al., 2013***) alignment software, and subsequently assigned to unique genes using HTseq software v0.10.024 (***Anders et al., 2015***). Subsequently, genes with fewer than 10 reads in fewer than 9 samples were excluded from further analysis. The resulting read counts matrix was normalized using the median-of-ratios method, implemented in the R package DESeq2 v1.42.1 (***Love et al., 2014***). Pairwise comparisons of differential gene expression were conducted across developmental stages using DESeq2. Genes were classified as differentially expressed if they exhibited a p-value less than 0.05 and a log2 fold change greater than 1 in absolute values.

### Integration analysis of DNA methylation and chromatin accessibility

An outside-in approach was employed to conduct a comprehensive analysis of DNA methylation and chromatin accessibility. To this end, the dataset from NCBI Gene Expression Omnibus (series GSE215396) (***Guerrero-Peña et al., 2023b***) of Assay for Transposase-Accessible Chromatin sequencing (ATAC-seq) corresponding to four developmental stages in whole turbot was employed (***Guerrero-Peña et al., 2023a***). Initially, two datasets were created: one of differentially methylated regions (DMRs) among all samples and another one of chromatin open peaks at all four stages. Subsequently, the intersectBed utility from Bedtools v2.28.0 package was employed to ascertain the regions of overlap. Only DMRs that intersected with the total chromatin open peaks by a distance of at least 100 bp were retained for further analysis. The mean methylation levels (average mCG/CG) of DMRs in an open chromatin region (OCR) were estimated using the DMLtest function from the DSS R package. The mean methylation levels for the three metamorphic stages were plotted using the heatmap.2 function from the gplots R package v3.13.1 (Spearman’s correlation matrix and Mcquitty clustering). Hypermethylated regions exclusively present at the climax of metamorphosis were explored in the ATAC-seq dataset. Among them, only those overlapping with differentially more open chromatin region at the same stage were extracted and used for further analysis.

### Genomic Features Annotation

A query gene model was constructed from the turbot genome annotation using the function *makeTxDbFromGFF* from the GenomicFeatures R package v1.42.3. Based on the query gene model, a breakdown of the genome was performed, with the following priority order: promoters (upstream = 2kb; downstream = 2kb), exons, introns and intergenic regions. Identified DMRs were assigned to these annotated genomic features from the turbot reference genome. The function *calcExpectedPartitions* from the GenomicDistributions R package was then performed to calculate the enrichment of DMRs for different genomic features. Chi-square tests were calculated for observed/expected overlaps on DMRs for each comparison group.

### CpG Island Annotation

Seven different reference genomes were downloaded from NCBI using the datasets command-line tool: human (GCF_000001405.29), mouse (GCF_000001635.9) and xenopus (GCF_ 000004195.4). Additionally, the following reference genomes were downloaded: zebrafish (GCF_000002035.4), lamprey (GCF_010993605.1), halibut (GCF_009819705.1), flounder (GCF_036983055.1), solea (GCF_019176455.1) and turbot (GCF_022379125.1). The CpG islands (CGI) of the entire reference genomes were predicted using two distinct algorithms. First, we evaluated the CGI content using Garden-Gardiner and Frommer’s (***Gardiner-Garden and Frommer, 1987***) algorithm. The following criteria were used: (1) A GC content of 50% or greater; (2) a length greater than 200 pb; (3) a ratio of observed/expected greater than 0.6. The observed/expected ratio of CpG dinucleotides in a given island can be calculated using the following formula: *Obs/Exp CpG = (Number of CpG · N)/(Number of C · Number of G)*, where the expected number of CpGs in an island is calculated as *Expected = (Number of C · Number of G)/(Length)*. We used the *cpg_lh* C program by LaDeana Hillier and UCSC based on the original work of Gos Miklem from the Sanger Center. Next, we applied Takai and Jones’s algorithm (***Takai and Jones, 2002***) for the same reference genomes with the following criteria: (1) GC content equal to or greater than 55%; (2) length greater than 500pb; (3) observed CpG/expected CpG of 0.65; where GC = (number G + number C)/(Length); *Observed CpG = (number CpG)/(Length)*; and *Expected = (GC/2)*^*2*^ (***Saxonov et al., 2006***). TJ’s algorithm was performed with the TaJoCGI software by Lucas Nell (***Nell, 2020***).

### Correlation of DNA Methylation and Gene Expression

In order to gain insight into the mechanisms of DNA methylation in turbot, we conducted a study to investigate the correlation between DNA methylation and gene expression in different genomic regions across all our samples. First, each genomic feature coordinate was retrieved from the reference genome using GenomicFeatures R package, and classified into the following categories: promoter, first intron, first exon, remaining introns, and remaining exons. The term “promoter” was defined as a region spanning 5 kilobases (kb) upstream and 1 kb downstream from the transcription start site (TSS). All annotated turbot genes were divided into ten expression deciles, with the first group comprising genes with the lowest level of expression and the tenth group including genes with the highest level of expression. Methylation levels were calculated for each genomic feature. The percentage of methylcytosine (mCG) to cytosine (CG) was calculated as follows: (average C methylated in the region) / (total number of C in the region) · 100. To obtain the gene IDs from the promoters and correlate them with gene expression, the AnnotationDbi R package was employed. A Spearman correlation was conducted between the methylation level (%mCG/CG) and gene expression for each genomic feature, employing the *cor*.*test* function from the *stats* R package. Genes belonging to the first three and last three deciles of promoters, first exons and first introns were extracted for further analysis.

### Mapping of DMRs to Genes and GO Enrichment Analysis

The turbot gene annotation was read with the function readTranscriptFeatures from the genomation v1.34.0 R package to map genes near the DMRs. The genes situated in relation to the genomic coordinates of interest were calculated with the function annotateWithGeneParts, with due consideration given to the nearest gene in accordance with the distance from the feature. The functional profiles of each gene cluster were computed using the compareCluster function from the ClusterProfiler R package v3.14.3 (***Yu et al., 2012***) to achieve GO enrichment analysis. An ad hoc turbot genome functional annotation, created with Sma3s (***Muñoz-Mérida et al., 2014***; ***Casimiro-Soriguer et al., 2017***), was employed as a custom background.

## Acknowledgments

This research was funded by the MCIN/AEI/10.13039/501100011033 grant number AGL2017-89648P to JR and by “ERDF A way of making Europe”. L. Guerrero-Peña was supported by a pre-doctoral fellowship of the Spanish FPI Research Training Program funded by Spanish Economy and Competitiveness Ministry, grant number (PRE2018-085475).

This work was made using a template from *eLife* journal provided under the CC BY 4.0 license with some minor style changes.

